# Retinal axial motion analysis and implications for real-time correction in human retinal imaging

**DOI:** 10.1101/2022.01.07.475424

**Authors:** Yao Cai, Kate Grieve, Pedro Mecê

## Abstract

High-resolution ophthalmic imaging devices including spectral-domain and full-field optical coherence tomography (SDOCT and FFOCT) are adversely affected by the presence of continuous involuntary retinal axial motion. Here, we thoroughly quantify and characterize retinal axial motion with both high temporal resolution (200,000 A-scans/s) and high axial resolution (4.5 µm), recorded over a typical data acquisition duration of 3 s with an SDOCT device over 14 subjects. We demonstrate that although breath-holding can help decrease large-and-slow drifts, it increases small-and-fast fluctuations, which is not ideal when motion compensation is desired. Finally, by simulating the action of an axial motion stabilization control loop, we show that a loop rate of 1.2 kHz is ideal to achieve 100% robust clinical in-vivo retinal imaging.

## 1. INTRODUCTION

A major challenge for *in vivo* retinal high resolution ophthalmic imaging with optical coherence tomography (OCT) is posed by artefacts caused by constant involuntary axial eye motion, from sources including head and body motion, physiological phenomena such as cardiac cycle, blood flow, pulsation, muscular and/or respiratory activity [1, 2]. For simple B-scan acquisition with SD-OCT devices, motion related artifacts can be automatically corrected during post-processing, but with the growing popularity of imaging methods based on volumetric SDOCT acquisitions with *en face* image reconstruction such as motion contrast-based OCT-Angiography (OCTA) [3], axial eye motion may provoke enough decorrelation to cause the false positive appearance of flow, significantly degrade the image quality, and add distracting artifacts in reconstructed *en face* slices [4].

Axial eye motion is also an important consideration for camera-based OCT techniques which acquire directly in the *en face* direction, such as time-domain full-field optical coherence tomography (FFOCT). FFOCT is an imaging modality capable of recording high-speed en-face sections of a sample at a given depth [5]. One very attractive characteristic of FFOCT is the use of spatially incoherent illumination to make the FFOCT lateral resolution robust to symmetric optical aberrations [6], which are the dominant aberrations in the eye (around 90%) [7]. By exploiting this phenomenon, we were able to apply FFOCT for high 3D resolution imaging of the living human retina. This achievement was possible owing to recent advances in FFOCT, including shaping the temporal coherence gate to match the retina curvature, achieving a wide field-of-view (5°*×* 5°) [8], stabilizing the axial retinal motion in real-time [9], and increasing signal-to-noise ratio (SNR) using the adaptive-glasses approach for ocular aberration correction [10]. Given its relative simplicity and small footprint, FFOCT holds promise for adoption by clinicians [11].

When acquiring a single en-face image in FFOCT, a micrometer stabilization precision can be obtained. However, longer acquisition durations on the order of a few seconds may be necessary in the clinical setting in order to allow image averaging to improve SNR. Therefore, it is necessary to accurately compensate for axial motion during image acquisition by moving the reference or the sample arm of FFOCT accordingly [9]. A few methods exist, but they are either used in conventional OCT, thus not suitable for FFOCT en-face imaging [12, 13], or are not sufficiently stable to remain within the coherence gate for acquisition durations over several seconds [9]. Indeed, our previous work [9] achieved an RMS error of around 10 µm, i.e. greater than the coherence gate width of 8 µm, which was nevertheless sufficient for a satisfactory 60 % imaging success rate in subjects with good fixation. However, for imaging in patients who may have poorer fixation, and eventual extension to *in vivo* functional imaging with dynamic FFOCT [14], a greater precision of axial tracking is desirable. Although the causes of axial retinal motion are well understood, the literature on axial retinal motion is still incomplete, as collected data had low temporal and axial resolution [1], and tested relatively few subjects [1, 9, 13], so that general conclusions on the most suitable design of a retinal axial motion stabilization control-loop could not be drawn.

In this Letter, we present the first characterization of the axial retinal motion with both high temporal resolution (200,000 A-scans/s) and high axial resolution (4.5 µm in water), over a typical data acquisition duration of 3 s. Characterization is performed on a 14-subject population, making it possible to come up with a statistical description reflecting the inter-subject variability. We also evaluate for the first time the direct influence of breathing on axial motion in our population, as this is a parameter that is clinically feasible to control during short image acquisition durations and which may affect stability. Finally, we draw conclusions on design considerations of axial retinal motion stabilization for robust clinical *in-vivo* retinal OCTA and FFOCT imaging.

## 2. METHODS

To measure the retinal axial position, we used a commercial spectral domain OCT system (SD-OCT; GAN611, Thorlabs). The SD-OCT system comprises a broadband superluminescent diode with 930 nm central wavelength and 100 nm bandwidth, providing a theoretical axial resolution of 4 µm in water. It presents an A-scan rate up to 248 kHz with a sensitivity of 84 dB and 1024 axial pixels, comprising an axial range of 2.2 mm in water. The galvanometer scanners of the SD-OCT were conjugated to the eye’s pupil with a 4-f system. The SD-OCT light beam arrives in the eye’s pupil with a 4 mm diameter.

To obtain statistics on axial retinal motion, 14 healthy subjects (age 22-80 years) free of ocular disease were invited to participate in the study. Research procedures followed the tenets of the Declaration of Helsinki. Prior to data collection, the nature and possible consequences of the study were explained, and informed consent was obtained from all subjects. This study was authorized by the appropriate ethics review boards (CPP and ANSM (IDRCB number: 2019-A00942-55)). Subjects were seated in front of the system and stabilized with a chin and forehead rest, without any pupil dilation nor cycloplegia. Image acquisition was realized in a dark room, maximizing the pupil dilation. They were asked to gaze at a yellow fixation cross displayed on a black screen using an LCD projector, maintaining fixation for about 4 seconds until acquisition was complete. To provide a good trade-off between acquisition speed and SNR, we chose to scan 1° field of view (FOV) of the retina with 256 A-scans at 200 kHz A-scan rate, corresponding to 1.28-ms exposure time for a single B-scan. With a 2-ms fly-back time, we were able to achieve a B-scan rate of 300 Hz. During image acquisition, the output power measured at the eye pupil plane was 850 µW, which is below the ocular safety limits established by the ISO standards for group 1 devices. To study the influence of breathing on axial retinal motion, two groups of data were obtained from our 14-subject population: while subjects held their breath or while they were breathing normally. The participants would indicate if they succeeded in holding their breath during acquisition. In the case of ambiguous data, a new dataset would be acquired.

After image acquisition, SD-OCT B-scan images were used to measure the retinal axial position using a previously introduced algorithm based on normalized cross-correlation [9]. Blinks were automatically detected by an intensity-based algorithm [9]. We extracted blink-free B-scan sequences, resulting in 3-s time series of retinal axial position data for all 14 subjects. Temporal power spectral densities (PSD) were calculated by a FFT routine.

## 3. RESULTS

### A. Temporal fluctuation of retinal axial position

Figure 1 (a-d) presents four typical examples of retinal axial position as a function of time. As exemplified here, slow drifts and relative fast oscillations could be observed in all 14 subjects. Still, the slow and fast motion amplitude and frequency varied among subjects. Figure 1 (e-h) present the corresponding axial motion power spectra density (PSD) for the same subjects in Figs. 1 (a-d). All PSDs present similar behaviour, with an energy decay in a *f* ^−*p*^ power-law, where f is the frequency, before reaching the noise-level plateau around 150 Hz. Note that the presence of pronounced peaks between 1-4Hz varied in the population.

**Fig. 1.**
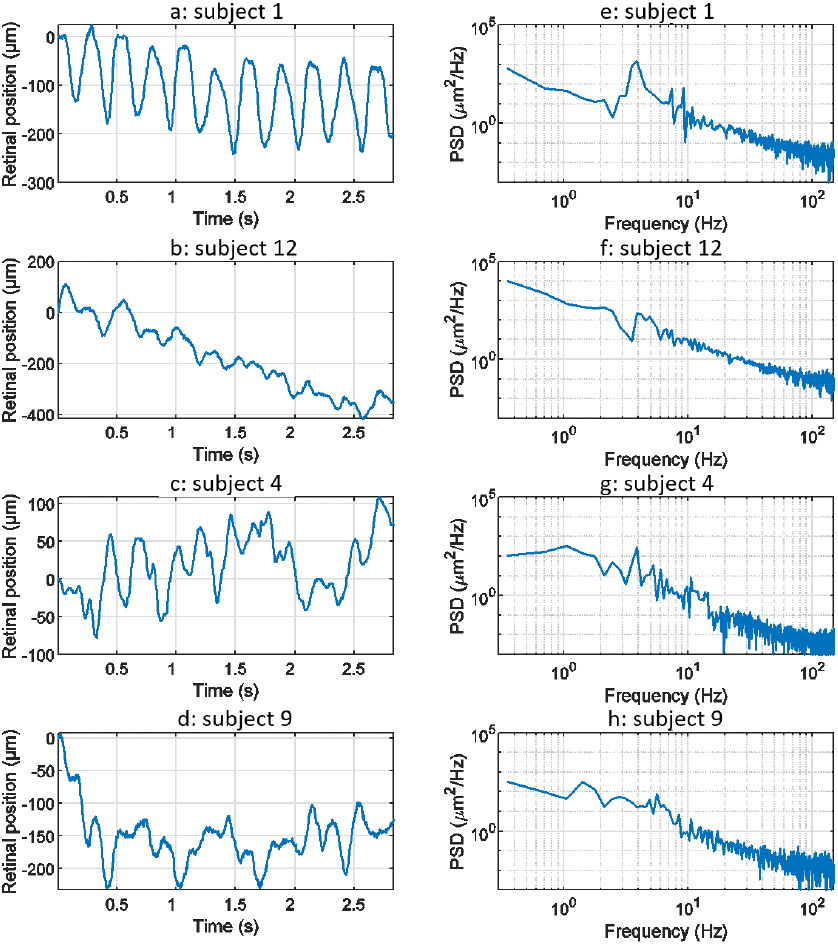
Dynamics of retinal axial position. (a-d) Four examples of the axial position temporal evolution over 3-s time series without blinks. (e-h) Corresponding temporal PSD computed for the retinal axial position. Note that subjects 1 and 12 held their breath, while subjects 4 and 9 breathed normally.

To draw statistics of retinal axial motion, and to investigate the effect of breathing on motion, we computed the standard deviation of each axial position time series for each of 14 subjects. Figure 2 shows temporal fluctuation of axial position under breath holding (blue bars) and normal breathing (red bars) conditions. Note that 9 subjects out of 14 presented a lower temporal fluctuation while holding their breath. Greater variations observed in subjects 12 and 13 are caused by larger drifts, most probably due to head motion. Two participants (subjects 1 and 6) were trained for eye fixation tasks. In contrast to retinal lateral motion, where experience in fixation can play an important role in the temporal fluctuation [15, 16], here no significant difference was noted, apart from a decrease of slow-and-large drifts linked to breathing and head motion.

**Fig. 2.**
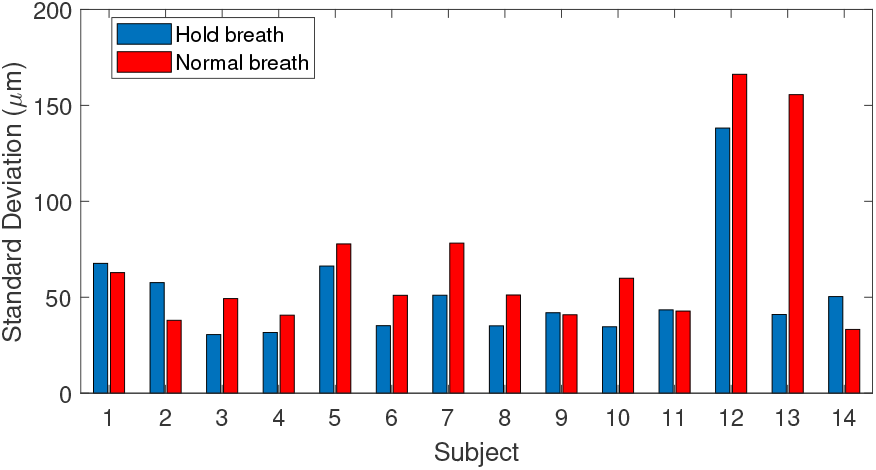
Fluctuations of axial position for 14 subjects tested during breath-holding (blue) and normal breathing (red), calculated as the standard deviation of axial position.

We computed the median ± standard deviation over the population of peak-to-valley (PV) amplitude, mean deviation from zero, root mean square (RMS) value and speed for both breath-holding and normal breathing conditions. Results of this statistical analysis is summarized in Table 1. During normal breathing, retinal axial motion presents higher PV amplitude, meaning that slow drifts are more important compared with the breath-holding condition. Indeed, it has been shown that breathing may cause slow and large drifts due to the movement of the subject’s head [1]. Not surprisingly, after filtering out slow drifts corresponding to less than 2 Hz temporal frequency, we found similar PV amplitude value for both conditions, 158 ± 52 µm for normal breathing compared to 160 ± 46 µm for breath-holding. Similar values of mean deviation from zero (26±15 against 25±11 µm) and RMS (32±18 against 31±12 µm) were also obtained for normal breathing and breath-holding conditions respectively. These results suggest that respiration induces axial motion at temporal frequencies up to 2 Hz, whereas respiration rate is around 0.2Hz [1]. On the other hand, the breath-holding condition exhibited a faster axial movement of 988 ± 209 µm/s, compared to 941 ± 194 µm/s in the case of normal breathing, for which the causes are still unclear. In both cases, computed speeds are in line with 958 µm/s median speed previously reported [2], and are assumed to be related to the systolic phase of the heart pulse.

**Table 1.** Axial motion statistics (median ± standard deviation) over the 14-subject population. Statistics were drawn in both normal breathing and breath-holding conditions in time series of 3-s duration. PV = peak-to-valley amplitude.

| | PV ($\mu\text{m}$) | Mean deviation ($\mu\text{m}$) | RMS ($\mu\text{m}$) | Speed ($\mu\text{m/s}$) |
| --- | --- | --- | --- | --- |
| Normal breathing | $276 \pm 154$ | $57 \pm 36$ | $68 \pm 42$ | $941 \pm 194$ |
| Holding breath | $230 \pm 98$ | $42 \pm 24$ | $52 \pm 28$ | $988 \pm 209$ |

Figure 3 presents the averaged PSD over all 14 subjects in both breath-holding and normal breathing conditions. The averaged PSD has a *f*^−2.2^ mean power-law before reaching the experimental noise-level at 150 Hz. Temporal frequencies below 1Hz present a factor of two lower PSD in the breath-holding condition than the normal breathing condition, highlighting the influence of respiration on axial motion. Remarkably, this difference is more pronounced at temporal frequencies associated to heartbeat (1-2Hz), showing a link between breathing and heartbeat on retinal axial motion. Indeed, apnea can induce a reduction in cardiac output and peripheral vasoconstriction [17, 18]. Both normal breathing and breath-holding conditions presented a similar behaviour in the range of 3-6 Hz, with some marked peak rises around 3-4Hz, which may be a high order harmonic of heartbeat rate [1] or related to pulsatile blood flow in the retina or in the choroid [19]. Meanwhile, similar to the computed speed behaviour, the PSD in the range of 7-150 Hz is more important in the breath-holding condition than under normal breathing.

**Fig. 3.**
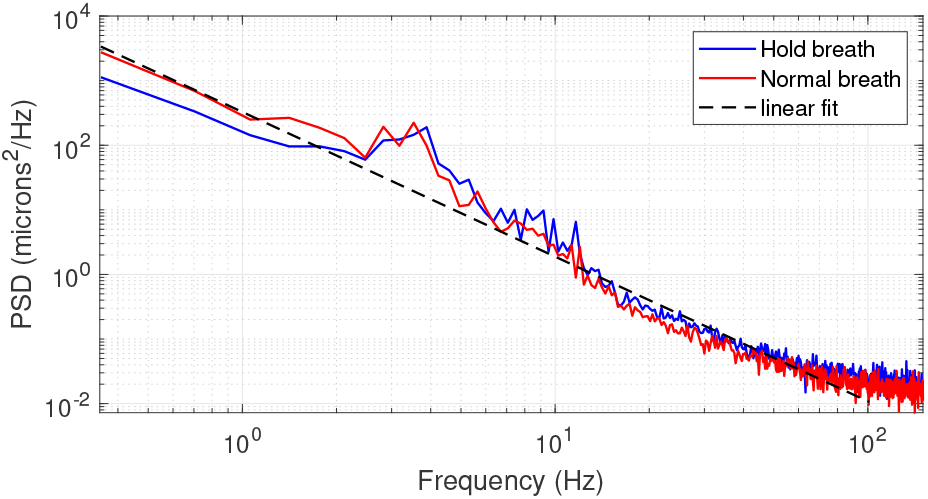
Averaged PSD over 14 subjects in breath-holding (blue trace) and normal breathing (red trace) conditions. The black dashed line gives a linear regression of the PSD following a *f* ^−2.2^ behaviour.

### B. Axial tracking loop simulation

Having characterized retinal axial motion, and knowing that breathing, heartbeat and pulsatile blood flow each play an important role, we next addressed the question of how fast the stabilization control-loop should run to precisely compensate for retinal axial motion in real-time for *in vivo* retinal imaging. To answer this question, we simulated the action of a retinal axial motion stabilization loop and integrator-based control scheme with a 0.5 gain (ensuring 45°stability margin) [20], assuming a two-frame delay [21]. Figure 4 displays the overall distribution of axial RMS tracking errors among all participants and for different stabilization control loop rates. This result provides a valuable tool for selecting the acquisition frequency for retinal axial motion measurement, and consequently the control-loop rate for stabilization. For instance, it indicates that at 150 Hz loop-rate, the residual axial motion should be approximately 10-µm rms in a normal breathing condition.

**Fig. 4.**
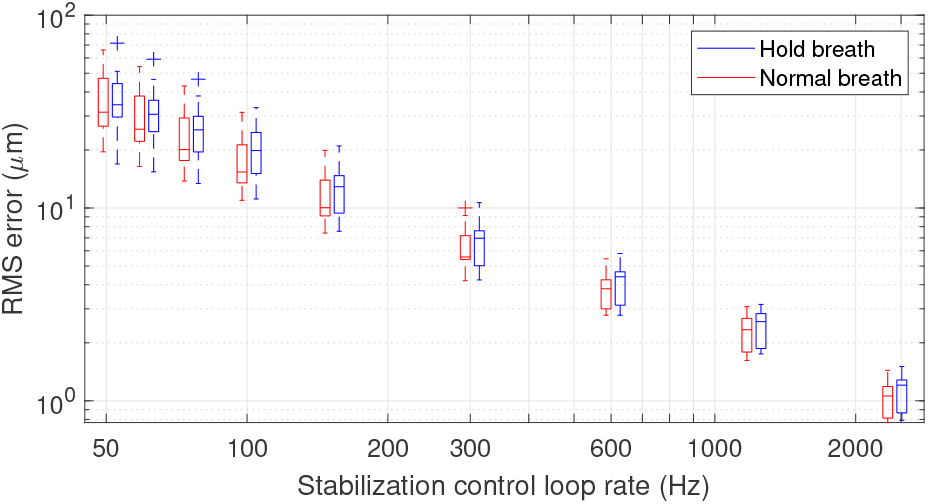
Box plot representation of residual RMS motion for different acquisition frequencies. The star (*) within the box identifies the median. The blue and red traces indicate median RMS tracking errors during breath-holding and normal breathing respectively. The length of the box represents the interquartile range (IQR), with the edges being the 25th (lower quartile) and 75th (upper quartile) percentiles. The ends of the whiskers represent minimum and maximum values. Values more than 1.5 IQRs are labeled as outliers (+). Note that the figure is plotted in a logarithmic scale.

In the context of applying dynamic (D-) FFOCT *in vivo*, the use of singular value decomposition (SVD) processing was recently proposed to filter out the axial displacement of the sample from the local fluctuations linked to intracellular motility [22]. This study showed that axial motion suppression works as long as residual axial motion after correction is smaller than the temporal coherence gate width [22, 23]. In FFOCT high-resolution *in vivo* retinal imaging, a temporal coherence gate of 8 µm is used [10]. Therefore, if we approximate the temporal fluctuation of retinal axial motion as a Gaussian distribution, one can establish that an RMS error below 4 µm will ensure a residual axial motion of amplitude lower than the 8 µm coherence gate width more than 95% of the time. An RMS error of 4 µm can be achieved on average with a stabilization control-loop rate of 400 Hz under normal breathing conditions. The same precision can be obtained under the breath-holding condition, but with a higher loop rate, here around 600 Hz, showing that holding breath does not necessarily help axial motion stabilization but may in fact hinder it. Indeed, although breath holding decreases low temporal frequencies, these are the frequencies that are better managed by an integrator-based control scheme [9, 20]. On the other hand, breath holding amplifies high temporal frequencies, which are also amplified by an integrator-based control scheme [9, 20].

In clinical application of FFOCT, one needs to consider the temporal fluctuation variability among subjects and ensure that the stabilization precision is enough to achieve robust imaging for 100% of the patients. Figure 5 highlights the proportion of subjects (of the 14 in total) where stabilization precision below 4 µm was obtained throughout the entire time series (i.e. 3-s) for different stabilization control-loop rates. To ensure robust FFOCT acquisition in all subjects, a control-loop rate of at least 1.2 kHz is necessary. Enticingly, this level of stability would also theoretically allow the first *in vivo* acquisition of dynamic FFOCT datasets, as axial motion would remain within the coherence gate width over the 3-s acquisition time required for a dynamic FFOCT image stack to probe temporal frequencies carrying intracellular dynamic signal (typically between 1-10Hz) [14]. This finding further supports the work of Pircher *et al* [13] who found that increased high loop rate of 1 kHz enabled the reduction of the residual axial motion below 5 µm in transverse-scanning OCT. As commonly done in OCTA, the reduction of the axial resolution can help to decrease the sensitivity to axial motion, thus relaxing the control loop rate [4]. For instance, a 600 Hz loop rate would be sufficient to ensure a residual RMS motion less than 8 µm in 100% of the patients for a 16-µm axial resolution (Fig.5). Another strategy to alleviate the control loop rate is the use of Linear-quadratic-Gaussian (LQG) control law [24]. Indeed, such an LQG approach could incorporate in the control strategy the specific spatio-temporal statistical structure of the axial retinal motion. Correcting the axial motion in real-time in volumetric SDOCT and OCTA en-face projections can also present advantages such as artifact suppression, optimization of SNR - given that the sensitivity decay with depth position would be avoided - access to the highest axial resolution, and decrease in the computational complexity of 3D registration algorithms.

**Fig. 5.**
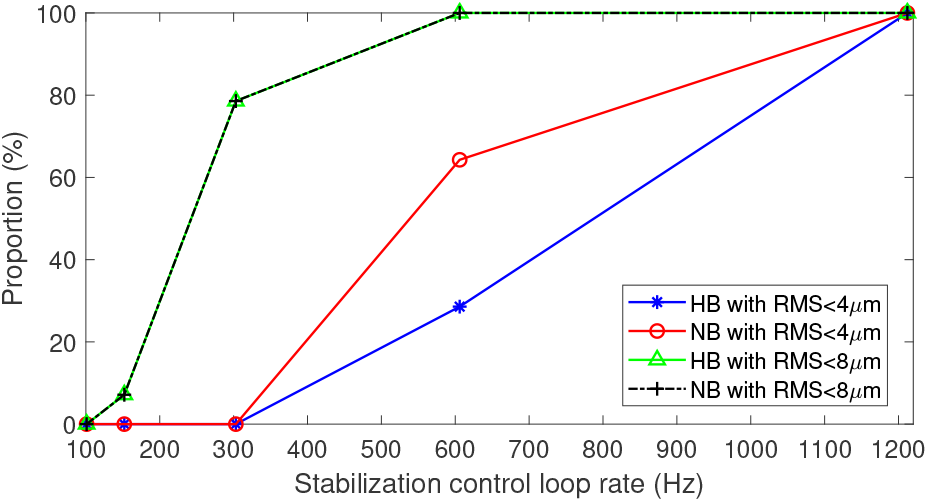
Proportion of data with residual RMS motion below 4 µm with respect to 8-µm axial resolution for *in vivo* FFOCT imaging, and below 8 µm with respect to 16-µm axial resolution for different acquisition frequencies. Blue trace for breath-holding and red trace for normal breathing.

## 4. CONCLUSION

In conclusion, this study has been one of the first attempts to thoroughly quantify and characterize retinal axial motion with high temporal resolution. The results of this experiment show that head motion during breathing, heartbeat and pulsatile blood flow significantly account for involuntary retinal axial displacement in the low-frequency domain below 5 Hz. We also demonstrated that breath holding can help decrease large-and-slow drifts. However, contrary to expectations, when holding breath, axial motion presented a higher speed and higher PSD for temporal frequencies from 7Hz to 150 Hz, which is not ideal when motion compensation is desired. Further investigation is necessary to understand the mechanism behind this phenomenon. Finally, by simulating the action of an axial motion stabilization control loop, we showed that a loop rate of 1.2 kHz is ideal to achieve 100% robust FFOCT in *in vivo* human retina, and could even pave the way to acquiring dynamic FFOCT *in vivo*. To relax the loop-rate constraint, a practical solution can be the combination of lower axial resolution and LQG control law.

## Acknowledgments

The authors thank all participants for their support, and Olivier Thouvenin and Yan Liu for fruitful discussions.

